# scTAM-seq enables targeted high-confidence analysis of DNA methylation in single cells

**DOI:** 10.1101/2022.04.11.487648

**Authors:** Agostina Bianchi, Michael Scherer, Roser Zaurin, Kimberly Quililan, Lars Velten, Renée Beekman

## Abstract

Single-cell DNA methylation profiling currently suffers from excessive noise and/or limited cellular throughput. We developed scTAM-seq, a targeted bisulfite-free method for profiling up to 650 CpGs in up to 10,000 cells per experiment, with a dropout rate of less than seven percent. scTAM-seq focuses sequencing coverage on informative, variably methylated CpGs that in many tissues and cell types make up a minor fraction of all CpGs. By applying scTAM-seq to B cells from blood and bone marrow, we demonstrate that it can resolve DNA methylation dynamics across B-cell differentiation at unprecedented resolution, identifying intermediate differentiation states that were previously masked. scTAM-seq additionally queries surface protein expression and somatic mutations, thus enabling integration of single-cell DNA methylation information with cell atlas data, and opening applications in tumor profiling. In summary, scTAM-seq is the first high-throughput, high-confidence method for analyzing DNA methylation at single-CpG resolution across thousands of single cells.

## Background

DNA methylation (DNAm) at CpG dinucleotides is an epigenetic mark extensively modulated in health and disease. DNAm has primarily been investigated in bulk samples, hindering the study of rare cell types, differentiation processes, and cellular heterogeneity. Single-cell DNAm (scDNAm) methods can overcome this limitation but currently require prohibitive sequencing efforts to cover the 28 million CpGs in the human genome. Available techniques with a cellular throughput of more than 100 cells produce very sparse datasets, where only 1-7% of the investigated CpGs are covered in a single cell [1–6]. Additionally, most CpGs in the genome are not informative to assess at single-cell level, as they are either constitutively (un)methylated, or display no variable methylation within the tissue of interest [7]. For instance, less than 4% of CpGs show a DNAm difference of more than 50% across the differentiation process of human B and T cells [8] (Suppl. Fig. 1). Moreover, bulk DNAm data is available for most human tissues and cell types [9], and can be used to identify CpGs with variable methylation [10]. Thus, targeted approaches that focus sequencing on variable CpGs can efficiently dissect intra-tissue DNAm heterogeneity and capture most of the variation observed in whole genome data (Suppl. Fig. 1), but have so far been limited to a throughput of less than 100 cells per experiment, and less than 60 investigated CpGs per cell [11–13].

**Figure 1.**
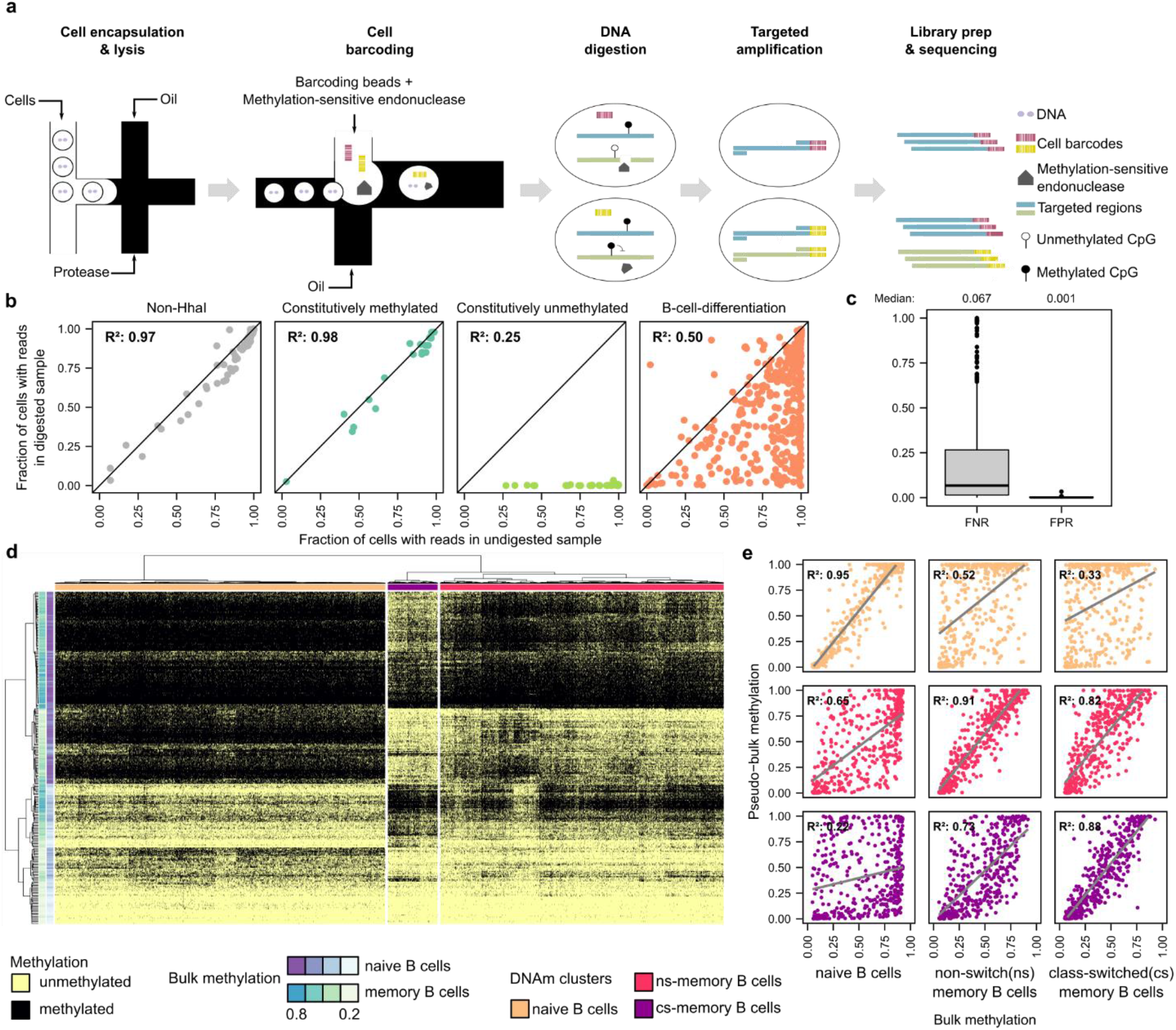
ScTAM-seq identifies cellular subtypes in peripheral blood B cells. **a**. Overview of the scTAM-seq workflow. **b**. Per amplicon comparison of fraction of cells with at least one sequencing read in the undigested and digested sample. **c**. False-negative rate (estimated on B-cell-differentiation amplicons, undigested control) and false-positive rate (estimated on the constitutively unmethylated amplicons, digested sample) across all amplicons of the respective class. **d**. Heatmap of single-cell DNAm values of 9,583 cells across 313 high-performance amplicons. **e**. Comparison of pseudo-bulk and bulk DNAm for the 424 B-cell-differentiation related amplicons. R^2^: Pearson’s correlation coefficient. The colors correspond to DNAm clusters in (**d**).

## Results and Discussion

We have developed scTAM-seq (single-cell Targeted Analysis of the Methylome) for measuring DNAm states at single-cell resolution in a high-throughput manner, with a median false-negative rate (FNR) of less than 7% and a median false-positive rate (FPR) of less than 0.2% at the level of single CpGs and single cells. To achieve this, we have combined and optimized single-cell PCR in droplets (Mission Bio Tapestri platform [14]) with the digestion of genomic DNA using a methylation-sensitive endonuclease. Among five candidate enzymes, we identified HhaI and SsiI as fully active in the Tapestri barcoding buffer (see Methods), and used HhaI in the following experiments. This enzyme selectively cuts unmethylated GCGC recognition sites, while leaving methylated sites intact. Hence, upon digestion of genomic DNA in barcoded single-cell droplets, amplicons containing targeted CpGs can only be amplified from methylated recognition sites (Fig. 1A). Based on the Tapestri platform, scTAM-seq enables the analysis of 650 CpGs in up to 10,000 cells, and can be combined with single-cell readouts of surface protein expression and somatic mutations [15, 16]. Thereby, it allows to characterize the surface phenotypes of cell types defined by DNA methylation, develop FACS schemes for newly identified cell types [17], and it has the potential to uncover clonal and sub-clonal development of tumors.

For a pilot study, we designed a panel of primers (Suppl. Tab. 1) to amplify 424 amplicons containing CpGs with dynamic DNAm during B-cell differentiation. These include CpGs differentially methylated among hematopoietic stem cells (HSCs), pre-B cells, immature B cells, naive B cells, and memory B cells [18]. In addition, as controls, we designed amplicons without HhaI recognition sites (non-HhaI) and amplicons covering CpGs that are constitutively methylated or unmethylated across B-cell differentiation. We applied scTAM-seq to B cells purified from either bone marrow or peripheral blood while simultaneously profiling the expression of 46 cell-surface proteins by staining cells with oligonucleotide-tagged antibodies [15, 16]. For both samples, we generated data from two experimental conditions, one digested by HhaI and one undigested control sample. Using the per-cell performance of the non-HhaI control amplicons as a quality filter, we obtained data for 5,340-9,583 cells per experiment (Methods and Suppl. Tab. 2).

To assess the FNR and FPR of scTAM-seq, we compared the performance between the digested and undigested bone marrow sample (Fig. 1B). Non-HhaI and constitutively methylated amplicons exhibited similar fractions of cells with mapped reads in both samples, showing an amplicon-dependent dropout rate. As this rate can be calculated from the undigested sample, it can be accounted for during downstream analysis of the digested sample (see Methods). Amplicons containing constitutively unmethylated CpGs in B cells only displayed reads in the undigested sample. From these amplicons, we inferred a median FPR (i.e., the number of cells with reads for a given constitutively unmethylated amplicon in the digested sample) of less than 0.2% (Fig. 1C), demonstrating high digestion efficacy. Finally, most of the amplicons targeting B-cell-differentiation CpGs showed a lower fraction of cells with reads in the digested sample, demonstrating selective digestion of unmethylated CpGs (Fig. 1B). For these amplicons, we calculated a median FNR (i.e., the number of cells without reads for a given B-cell-differentiation amplicon in the undigested control) of 6.7% at the level of single cells and single CpGs (Fig. 1C) [11, 12], which is 12-15-fold lower than current droplet- or split-pool methods [1–4]. We note that FNR is influenced by the GC content of the amplicon (Suppl. Fig. 2).

**Figure 2:**
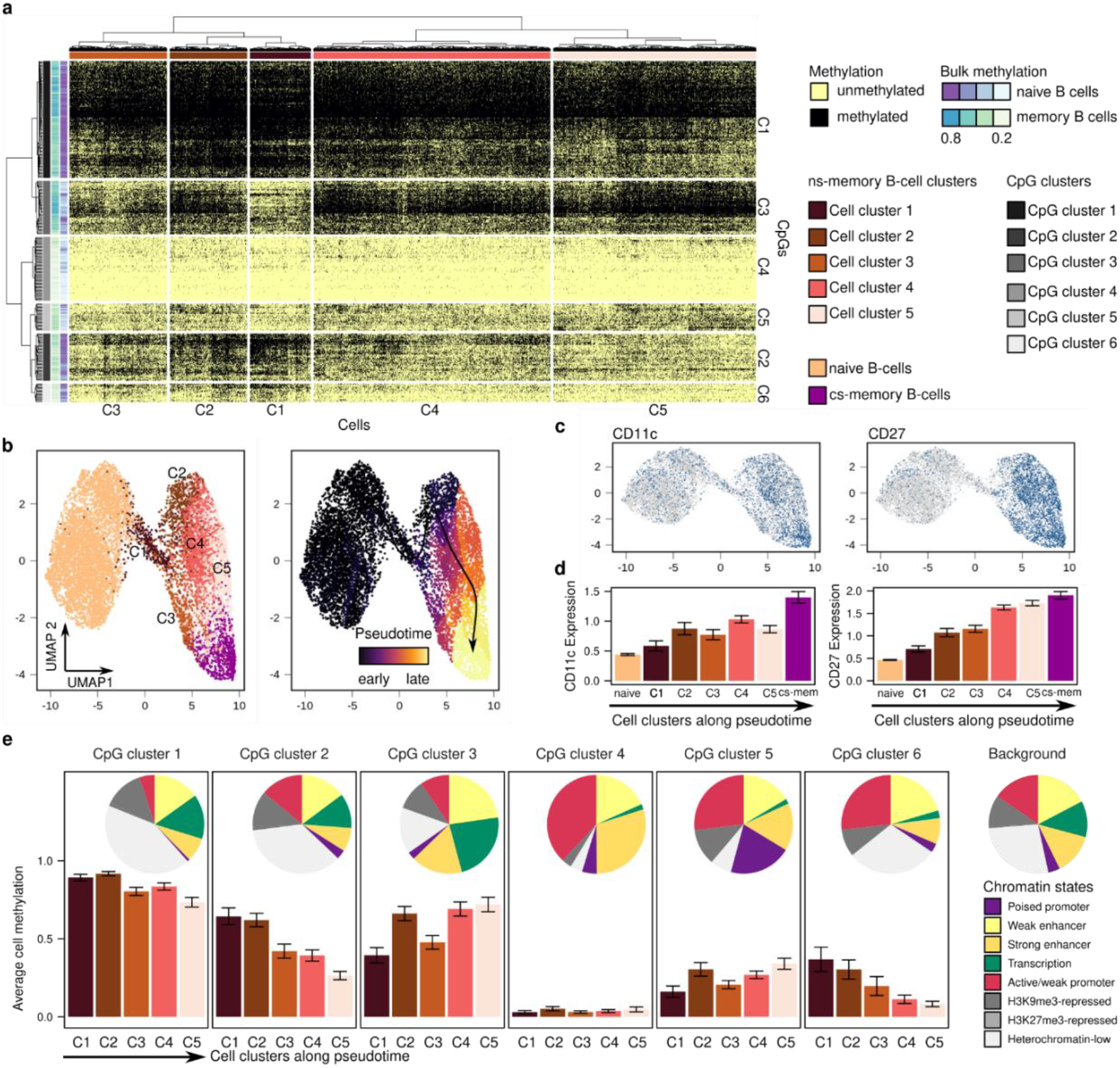
scTAM-seq identifies cellular states associated with proliferation. **a**. Heatmap showing the binarized, single-cell DNAm matrix for 4,100 ns-memory B cells in the 313 high-performance amplicons. Five clusters of ns-memory B-cells and six CpG clusters were defined based on a hierarchical clustering (binary distance, Ward’s method). **b**. Low-dimensional representation of the binarized data matrix for all cells (naive, cs- and ns-memory B cells) using UMAP. The pseudotime was inferred with Monocle3 [26]. **c**. Surface protein expression within the UMAP-space. The protein expression data was binarized using a cutoff of 1 for the CLR-normalized counts. **d**. Surface protein expression for the different clusters (ordered by increasing pseudotime) identified in **a**. Shown is the mean and two times the standard error within each of the clusters. **e**. Average DNA methylation value per CpG- and cell cluster were estimated by computing the fraction of all methylated amplicons in a given CpG and cell cluster. The error bar indicates two times the standard error across all cells of a cell cluster. The pie chart indicates the genomic distribution of the CpGs within each CpG cluster according to chromatin states of naive, germinal center, ns-, and cs-memory B cells defined in Beekman *et al*. [22].

To evaluate the ability of scTAM-seq to resolve cell populations, we first analyzed B cells from a peripheral blood sample [15, 16]. Unsupervised clustering of high confidence CpGs (FNR < 0.25) identified three clusters. We used reference bulk DNA methylation data to identify these clusters as naive, non-switched (ns-) memory and class-switched (cs-) memory B cells (Fig. 1D). Surface protein data further validated these cluster assignments and were in line with single-cell CITE-seq data [17] (Suppl. Fig. 3, 4). We next computed pseudobulk DNAm values per cell-type cluster for all CpGs included in the assay while accounting for the FNR per amplicon (Methods). We found a strong correlation between pseudobulk and reference bulk data [18, 19] of the corresponding cluster (R^2^ ≥ 0.88, Fig. 1E, Suppl. Fig. 5), confirming the high accuracy of scTAM-seq.

**Figure 3.**
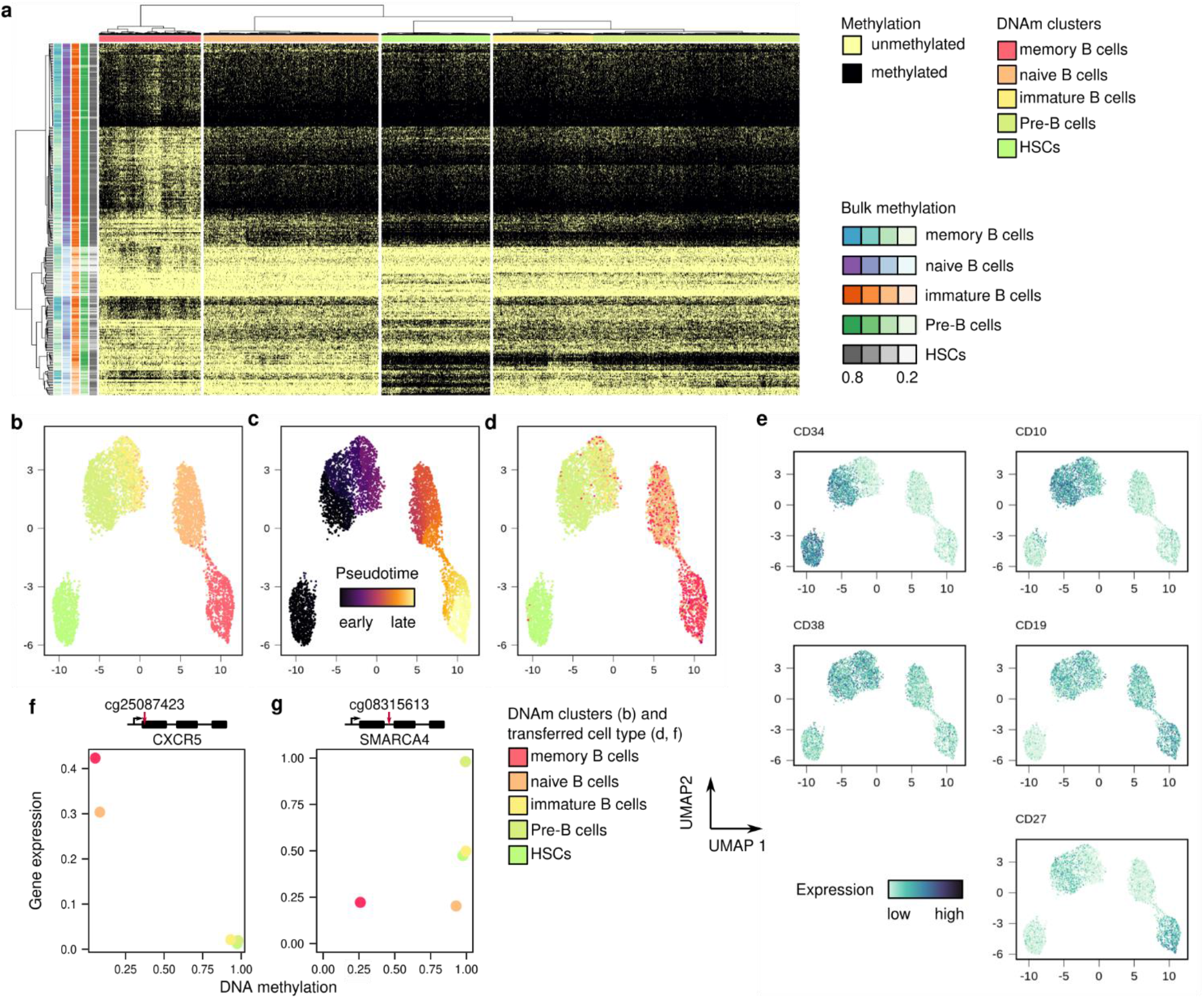
ScTAM-seq captures the B-cell differentiation process in bone marrow. **a**. Heatmap of single-cell DNAm values for 5,340 cells and 313 amplicons. **b-d**. Visualization of DNAm data in low-dimensional space (UMAP), with labels inferred from bulk DNAm (**b**), across differentiation pseudotime (inferred with Monocle3 [25], **c**), or with labels transferred from CITE-seq data (**d**). **e**. Visualization of surface protein expression for cells embedded in the DNAm UMAP. The value shows the CLR-normalized expression values (see Methods). **f, g**. Relationship between DNAm and log-normalized gene expression across B-cell differentiation showing a positive (**f**, CXCR5, promoter region) and negative correlation (**g**, SMARCA4, intronic region), respectively.

To evaluate scTAM-seq’s ability to resolve intra-population heterogeneity, we performed a focused analysis of the ns-memory B cells. We identified substantial heterogeneity within this population at the DNA methylation level that was linked to differences in the surface expression of CD27 and CD11c surface proteins (Fig. 2A-D). The gradual gain of CD27 appeared to be associated with gradual progression of cellular differentiation, with CD27-negative ns-memory B cells representing early, atypical memory B cells that are present at the highest frequencies at birth [20, 21]. Furthermore, linking the different DNA methylation patterns within the ns-memory B cell cluster to chromatin states [22] showed that one CpG cluster that loses methylation (CpG cluster 1) is enriched for heterochromatin (Fisher’s test p-value: 2.28×10^−9^), while gain of methylation (CpG cluster 5) occurs more frequently in polycomb-associated poised promoters (Fisher’s test p-value: 2.7×10^−4^) (Fig. 2E, F). These patterns have previously been linked to proliferative history [23], showing the gradual increase in proliferative history upon cellular differentiation. Additionally, subsets of memory B cells expressing CD11c have previously been described as more prone to differentiate to plasma cells, and are enriched in autoimmune diseases [24, 25]. Overall, our results show the potential of our method to resolve cell types to an unprecedented resolution, characterizing subpopulations masked until now at bulk DNA methylation level, and linking them to other mechanisms such as gradual differentiation and proliferative history.

We next investigated scTAM-seq’s potential to profile DNAm dynamics during B-cell differentiation in bone marrow. Dimensionality reduction and clustering using scDNAm data revealed five groups of cells arranged in a putative B-cell differentiation pseudotime trajectory [26]. We first labelled the clusters as stem- and progenitor cells, pro/pre-B cells, immature B cells, naive B cells, and memory B cells based on pseudotime and reference bulk DNAm patterns (Fig. 3A-C, Suppl. Fig. 5, 6). To identify cell types in an automated manner without making use of bulk DNAm levels, we transferred labels from a single-cell CITE-seq reference [17] using the surface protein data captured by scTAM-seq (Fig. 3D). These annotations, as well as the expression of specific surface markers (Fig. 3E), confirmed the order of B-cell differentiation stages by pseudotime analysis. Next, we demonstrated that the surface protein data can be used to integrate scDNAm and RNA-seq data into a common reference space (Suppl. Fig. 8). This allowed us to identify CpGs anti-correlated and correlated with gene expression throughout the differentiation trajectory, as exemplified for CXCR5 (negative correlation, gene promoter) and SMARCA4 (positive correlation, intronic region, Fig. 3F, G). Together, these analyses demonstrate the abilities of scTAM-seq to dissect DNA methylation heterogeneity of complex cellular populations.

As scTAM-seq is a targeted method, only up to 650 CpGs of the 28 milion CpGs in the human genome can be analysed. On the other hand, genome-wide approaches for scDNAm profiling largely suffer from limited cellular throughput and high data sparsity. Additionally, for most tissues and cell types, bulk DNAm data is available and only few CpGs are variable across cell differentiation (Suppl. Fig. 1). Thus, targeting only a few, highly variable CpGs is promising to reveal DNAm heterogeneity across cellular differentiation at unprecedented accuracy. The Mission Bio tapestri platform has recently been extended to cover 1,000 amplicons, which increases the coverage of scTAM-seq even further. The selection of potentially interesting CpGs for scTAM-seq is facilitated by tools for the identification of variably methylated sites from bulk data [10]. A pipeline automating the selection of target sites for scTAM-seq is available at https://github.com/veltenlab/CpGSelectionPipeline. Of note, the use of HhaI limits the analyzable CpGs to 1.76 million in the human genome, but further sites can be analyzed using other enzymes such as SsiI.

## Conclusions

In summary, scTAM-seq is a powerful method for investigating DNAm dynamics at single-cell and single-nucleotide resolution. Due to its targeted nature, it focuses sequencing coverage on CpGs variably methylated in a cell population of interest, alleviates data sparsity and thus enables high-throughput cell-to-cell comparisons at the level of single CpGs.

## Declarations

### Ethics approval and consent to participate

All experiments involving human samples were approved by the clinical research ethics committee of the Hospital Clinic of Barcelona (number HCB/2018/1046 and HCB/2019/0018) and were in accordance with the Declaration of Helsinki.

### Consent for publication

Not applicable

## Availability of data and materials

Raw and processed DNA methylation and surface protein expression data is available from the Gene Expression Omnibus (GEO) under accession number GSE198019. All associated code to reproduce the analysis is available from https://github.com/veltenlab/scTAM-seq-scripts and a pipeline for selecting CpGs for scTAM-seq from https://github.com/veltenlab/CpGSelectionPipeline.

## Authors’ contributions

L.V. and R.B. conceptualized the project with contributions by A.B., M.S. and R.Z., R.Z. and R.B. selected the restriction enzymes, R.Z. and K.Q. tested restriction enzyme efficiencies, A.B. and R.Z. generated scTAM-seq data, A.B., M.S., L.V. and R.B. designed the amplicon panel, A.B. and M.S. performed data analysis, M.S. generated the CpG selection pipeline, A.B., M.S., L.V. and R.B. wrote the manuscript. All authors commented on the manuscript.

## Acknowledgements

We thank Dr. Holger Heyn and Sara Ruiz Gil for technical support.

## Supplementary materials

- Supplementary Figures 1-10: See enclosed pdf file.
- Supplementary Table 1: Design overview of the panel of targeted regions.
- Supplementary Table 2: Sequencing details per samples and cells.
- Supplementary Table 3: Complete list of methylation-sensitive endonucleases potentially compatible with scTAM-seq.
- Supplementary Table 4: Non-HhaI amplicons used for cells selection.
- Supplementary Table 5: PCR program for DNA digestion and targeted amplification steps.
- Supplementary Table 6: Description of the doublet removal step for scTAM-seq.
- Supplementary Table 7: Correlation analysis between target CpGs and gene expression.

## METHODS

### Amplicon panel design

#### Enzyme selection

Using the REBASE database (http://rebase.neb.com/rebase/rebase.html [1]), 60 DNA-methylation-sensitive restriction enzymes, targeting 33 restriction sites, were selected fulfilling the following criteria: (i) digestion activity is completely blocked by DNA methylation (5mC), (ii) enzyme is sensitive to heat inactivation, (iii) recognition site does not contain Ns (representing any nucleotide), (iv) recognition site harbors a single CpG, (v) enzyme is commercially available. The complete selection of enzymes is available in Suppl. Tab. 3. Next, we chose four enzymes to test in the Tapestri Barcoding Mix buffer: AciI (NEB, recognition site CCGC), HpaII (NEB, recognition site CCGG), HpyCH4IV (NEB, recognition site ACGT), and HhaI (NEB, recognition site GCGC), with 3.7M, 2.3M, 2.2M, and 1.3M recognition sites in the human genome (hg38), respectively. To comply with criterium (iv), sequence contexts in which two CpGs occur, e.g., CCGCG for AciI and, CGCGC or GCGCG for HhaI, are not included within this list; but without this consideration, there are 4.2M AciI, and 1.76M HhaI recognition sites in the human genome. These numbers were calculated using Biostrings v2.56.0, R package (https://bioconductor.org/packages/Biostrings).

The activity of these four enzymes was tested as follows. Shortly, PCR products were generated from genomic DNA of Jurkat cells using the primers: 5’-TTCCACGTTTTTCTTTCATGC-3’ and 5’- GCAGTCGTTGGTTGGAAACT-3’ for AciI, 5’-CCCAGGCGTTTGTTAAAGAG-3’ and 5’- GCATGAAAGAAAAACGTGGAA-3’ for HpaII, 5’-TGGCTGTAGCCAGTTCTCAA-3’ and 5’- AAGGACACGCCTCTCACACT-3’ for HpyCH4IV, 5’-GGGGATCAATCACCATATGAA-3’ and 5’- TGGCTGATGGGATCAACAAT-3’ for HhaI, followed by digestion of the respective amplicons for 30 minutes at 37°C in the standard buffer provided by the company, or the Tapestri Barcoding Mix buffer. HpaII and HpyCH4IV showed only partial digestion in the Tapestri Barcoding Mix buffer. On the contrary, HhaI showed the same ability to digest unmethylated PCR products in both buffers (data not shown) and was chosen for the follow-up scTAM-seq design. AciI did not show complete digestion in the Tapestri Barcoding Mix buffer, but its isoschizomer SsiI (Thermo Fisher) showed almost complete digestion, comparable with the standard buffer (data not shown), representing an alternative enzyme compatible with scTAM-seq.

#### Target regions selection

Target regions were selected fulfilling the following criteria: (i) containing one HhaI recognition site in a 300 bp window (except for non-HhaI amplicons, which do not contain any recognition site), (ii) GC content between 45 and 65 percent. The first criterium reduces the number of CpGs that can be analyzed by SsiI (scTAM-seq compatible) and HhaI to 1.6M and 0.8M, respectively. Bulk DNA methylation data of B-cell subpopulations (progenitor cells, pre-BI and pre-BII cells, immature, naive, non-class switch and class-switched memory, plasmablasts, and plasma cells) obtained by 450K array analysis (Illumina) were mined from Kulis *et al*. [2]. Using this data, we identified 1,753 CpGs that show dynamic DNA methylation during B-cell differentiation that can be digested by HhaI and fulfill the criteria above. Next, to select the ∼450 CpGs with the highest predictive value for the B-cell populations of interest, single-cell DNA methylation datasets were simulated using the bulk DNA methylation values of the 1,753 CpGs (considering an allelic false-negative rate of 0.2 and an allelic false-positive rate of 0.1). This data was employed to run a regularized linear model (glmnet R package [3], parameters: λ = *e*^−4^, α = 0.9), leading to the selection of a panel of 428 CpGs. Furthermore, to improve the limited separation between early B-cell subpopulations, 32 CpGs showing differential methylation among these populations (pairwise comparisons, FDR <0.05, minimum DNA methylation difference 0.25) were added to the panel yielding a final selection of 451 CpGs. As controls, we selected 50 constitutively unmethylated CpGs within an HhaI recognition site (top 50 CpGs with lowest DNA methylation levels in B cells, bulk DNA methylation <0.06 in all samples) and 30 CpGs constitutively methylated within an HhaI recognition site (top 30 CpGs with highest methylation levels in B cells, bulk DNA methylation >0.94 in all samples), as well as 96 amplicons without HhaI recognition site.

#### Amplicon design

Using the Tapestri Designer tool (https://support.missionbio.com/hc/en-us/articles/4404329631895-Tapestri-Designer-User-Guide) amplicons were designed spanning our CpGs of interest. Some regions yielded no amplicons, leading to a final panel covering 424 B-cell-differentiation related CpGs, 21 constitutively methylated CpGs, 32 constitutively unmethylated CpGs, and 87 non-HhaI regions. A complete overview of the panel design can be found in Suppl. Tab. 1.

#### Recommendations for panel design

Amplicon performance is related to GC content, we recommend excluding amplicons with a GC content above 0.65 (Suppl. Fig. 2). Non-HhaI amplicons are essential to distinguish between cells and empty droplets after sequencing, we recommend including at least 50 amplicons of this type in the design. To aid the inclusion of these control regions in other panels, we provide the exact locations of the high-confidence non-HhaI amplicons used for cell selection in our study in Suppl. Tab. 4.

### Sample preparation

#### Sample description

Peripheral blood samples were obtained from two healthy donors (female age 57, male age 49) from the *Banc de Sang i Teixits* (Catalunya, Spain). Frozen Bone Marrow mononuclear cells were purchased from StemExpress® (Folsom, CA, USA, Cat. num. BMMNC050C; Lot. 2106150113) and correspond to a 22-year-old healthy male donor. All experiments involving human samples were approved by the clinical research ethics committee of the Hospital Clinic of Barcelona (number HCB/2018/1046 and HCB/2019/0018) and were in accordance with the Declaration of Helsinki.

#### Isolation of B-cell subpopulations from peripheral blood

Briefly, peripheral blood was collected and stored at room temperature (15-25°C). Within 24 hours after sample collection, B-cell subpopulations were isolated using the RosetteSeqTM Human B Cell Enrichment Cocktail (StemCellTM Technologies; Cat. num. 15024) followed by a Ficoll®-Paque Premium (Gmbh; Cat. num. 17-5442-02) density gradient centrifugation. B-cell subpopulations were cryopreserved in heat-inactivated Fetal Bovine Serum (GibcoTM; Cat. num. 10270106) supplemented with 10% DMSO, until the day of the experiment.

#### Flow cytometry cell sorting

B-cell subpopulations from PBMCs were stained with the monoclonal antibody anti-CD19(PE) (Clone HIB19; Invitrogen; Cat. num. 12-0199-41) in a 1:20 dilution for 30 minutes on ice. Bone marrow mononuclear cells were stained for 30 minutes on ice with the following antibodies: anti-CD19(APC/Cy7) (clone HIB19; Biolegend; Cat. num. 302217) in a 1:20 dilution, anti-CD38(APC) (clone HIT2; eBioscience; Cat. num. 17-0389-42) in a 1:30 dilution, anti-CD123(PE) (clone 763; BD; Cat. num. 561058) in a 1:50 dilution, anti-CD10(PE/Cy7) (clone Hi10a; Biolegend; Cat. num. 312213) in a dilution 1:20 and anti-CD34(AF488) (clone 581; Biolegend; Cat. num. 343517) in a dilution 1:100. All samples were sorted using the BD Influx and FACSAria II SORP cell sorters. A purity of 90-95% CD19+ cells was obtained after sorting PBMC samples. The bone marrow sample was sorted for the following subpopulations: (S1) CD24+/CD38-, (S2) CD38+/CD34+/CD10+/CD123- and (S3) CD19+/CD34-, resulting in 9.25% of S1, 6.5% of S2 and 84.25% of S3 when considering the total number of sorted cells. These subpopulations were mixed in a ratio of 10% of S1, 22.9% of S2, and 67.1% of S3 for a total of 1M cells.

#### Oligo-tagged antibody cell staining

Following sorting, cells were stained with oligo-tagged antibodies following the instruction in the Tapestri® Single-Cell DNA + Protein Sequencing User Guide V2 (Tapestri User Guide V2, https://support.missionbio.com/hc/en-us/articles/360062406493-Tapestri-Single-cell-DNA-Protein-Sequencing-V2-User-Guide). Briefly, 1M cells were stained with TotalSeq-D Heme Oncology Cocktail (BioLegend; Cat. num. 399906) reconstituted in 59 μl of Cell Staining buffer (BioLegend; Cat. num. 420201) and adding 1 μl of TotalSeq-D0154 anti-CD27 antibody (clone O323, BioLegend; Cat. num. 302861) for 30 minutes on ice.

### scTAM-seq and surface proteins libraries preparation and sequencing

#### Samples processing using the Tapestri instrument

A total of 120,000-140,000 cells were loaded into a Tapestri microfluidics cartridge. Upon encapsulation, cells were lysed. To obtain the digested samples, the methylation-sensitive restriction enzyme was added to the barcoding master mix as follows: 288 μl of Tapestri Barcoding Mix V2, 5 μl of highly concentrated HhaI enzyme (150,000 U/mL, NEB; Cat. Num. R0139B-HC1), while keeping the remaining reagents as stated in the Tapestri protocol (5 μl Forward Primer Pool, 2 μl Antibody Tag Primer). To ensure the activity of the enzyme, the PCR program was modified as shown in Suppl. Tab. 5, introducing a 30-minutes step at 37°C, prior to the targeted amplification. This step was introduced for the digested and undigested samples to avoid bias in the amplification performance. The DNA digestion and targeted PCR steps were performed in a T100 Thermal Cycler (Bio-Rad). All other sample processing steps were performed following the Tapestri User Guide V2. All Tapestri related reagents were obtained using Tapestri Single-Cell DNA Custom Kits and Cartridge (Mission Bio, Inc; Cat. num. MB02-0001 and MB03-0034).

#### scTAM-seq and surface protein library purification and sequencing

The Tapestri User Guide V2 was followed to prepare the libraries, with a few modifications listed here. Briefly, PCR products were retrieved from individual droplets, and purified with 0.7X Ampure XP beads (Beckman Coulter; Cat. num. A63881), to split the scTAM-seq library bound to the beads from the surface proteins library in the supernatant. Illumina i5/i7 sequencing indexes were added to the scTAM-seq PCR products, followed by two steps of purification, using 0.6X and 0.65X Ampure XP beads (these are different from the ratios indicated in the Tapestri protocol). PCR products from the surface proteins library were incubated with Tapestri Biotin Oligo at 96 °C for 5 minutes, followed by incubation on ice for 5 minutes, and purification using Tapestri Streptavidin Beads. Afterwards, the beads were used as PCR templates for the incorporation of i5/i7 Illumina indices, followed by purification using 0.9X Ampure XP beads. The quality of all scTAM-seq and surface proteins libraries were assessed by Bioanalyzer (Suppl. Fig. 9). Libraries were pooled and sequenced on the Illumina NovaSeq6000 platform at the CNAG-CRG Sequencing Unit, at a sequencing depth of 260 M reads/library in case of surface proteins libraries, and 420 M reads/library for scTAM-seq libraries (Suppl. Tab. 2).

### Bioinformatic analysis

#### Raw data processing

Raw sequencing data was processed using a customized version of the Mission Bio Tapestri Pipeline v2 (https://support.missionbio.com/hc/en-us/sections/360006255314-Tapestri-Pipeline). Briefly, cell barcodes were extracted from the raw FASTQ data files and sequencing adapters trimmed using cutadapt [4] v2.5. Trimmed reads were aligned to the reference genome version ‘hg19’ using bwa-mem [5] v0.7.12. Subsequently, read pair information was verified using PicardTools (v1.126, https://github.com/broadinstitute/picard), and quantified with samtools [6] (v1.9), and the cell barcode distribution was computed using the python scripts provided by Mission Bio. As a final step, we identified cells from the barcodes as follows: we exclusively used the control amplicons to determine barcodes that can reliably be called as ‘cells’ according to their overall read counts, and adapted the original cell detection method implemented by Mission Bio (https://support.missionbio.com/hc/en-us/articles/360042381634-Cell-calling) as follows. We only used those control amplicons that were reliably captured in most of the cells (Suppl. Tab. 4). To determine a read count cutoff on a per amplicon and per cell basis, we focused on the number of amplicons in the panel and computed the threshold as the minimum of 10 and 0.2 times the average number of reads for those cells covered by at least eight times the number of amplicons in the panel (see https://support.missionbio.com/hc/en-us/articles/360042381634-Cell-calling for a more detailed description). Then, we call as cells those barcodes having more than the determined threshold of reads in at least 70% of the amplicons. To determine potential doublets, we employed the DoubletDetection (http://doi.org/10.5281/zenodo.2678041) software, which removed a maximum of 16% of cells from our data (Suppl. Tab. 6, Suppl. Fig. 10). The pipeline has been implemented in *bash* and is available from the GitHub repository (https://github.com/veltenlab/scTAM-seq-scripts).

#### Clustering analysis and dimension reduction

To obtain cell type clusters, we first binarized the cells by amplicon DNA methylation matrix according to the presence of at least one sequencing read. Due to the low FPR computed in the control experiment, we found that a cutoff of one sequencing read reliably differentiated methylated from unmethylated CpGs for an individual cell. We selected high-performance amplicons as those amplicons that are present in at least 75% of the cells in the undigested control, leading to 313 amplicons selected from the control sample (bone marrow). We clustered this matrix using the binary distance (i.e., the fraction of methylation calls that are different between any two vectors divided by the fully methylated states) and Ward’s minimum variance method (‘ward.D2’ option in the ‘hclust’ R function). We selected three clusters for the peripheral blood and five clusters for the bone marrow data, since we observed a separation into previously defined cell types.

To obtain a low-dimensional representation of the data, we used the read count matrix obtained from scTAM-seq after removal of doublets and selected the 313 high-performance amplicons associated with B-cell differentiation. After binarizing as stated above, we normalized the data using Seurat v4.0 [7] with the functions ‘NormalizeData’ (‘LogNormalize’ method), and then executed ‘ScaleData’, ‘RunPCA’, ‘FindNeighbors’ (dimensions 1-11), and ‘RunUMAP’ using all of the features. The cells of the bone marrow sample were annotated using the information from bulk data into five cell types. Then, we used Signac (v1.2) [8] and Monocle3 (v1.0) [9] to infer a trajectory in the low-dimensional space (‘learn_graph’ and ‘order_cell’ functions).

For the protein data, we used Seurat v4.0 and used the centered log ratio (CLR)-method to normalize the data. All protein expression values shown in the paper represent the CLR-normalized expression values.

#### Computing pseudo-bulk DNA methylation values

We aimed at obtaining a pseudobulk DNA methylation level for all cells in a given cluster on a per-CpG basis correcting for the dropout rate observed in the undigested sample. Notably, we aimed at inferring pseudobulk methylation values for all 424 B-cell differentiation amplicons, also for those with elevated dropout rates. We defined the observables for amplicon *i* and cell cluster *c*, which can be computed directly from the experiments as follows:

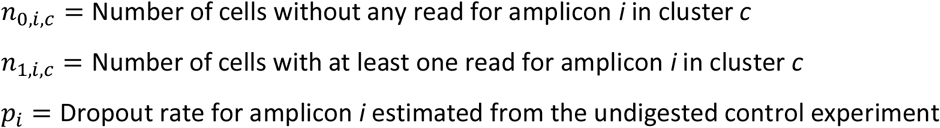

The parameter we aim to estimate is the DNA methylation value for cluster *c* in amplicon *i*:

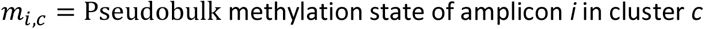

We assume that the reads that we obtained can be modelled using a Binomial distribution given the following formula:

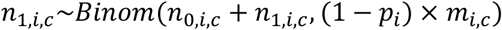

Additionally, we assume that we do not have prior information about the true underlying DNA methylation state and thus use an uninformative prior:

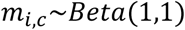

We use the Hamiltonian Monte Carlo algorithm as implemented in the ‘rstan’ R-package [10] to obtain the parameter *m*_*i,c*_ (pseudobulk methylation value) for each cluster separately.

#### Integrating scTAM-seq with scRNA-seq data

Since 33 of the 46 surface proteins are also measured in a single-cell proteo-transcriptomic atlas [11], we used this overlap to perform cell label transfer and data integration of the bone marrow data with the CITE-seq reference atlas [11] using Seurat v4.0 [7]. Multimodal nearest neighbors are defined between the antibody and transcriptomic modality in the reference atlas using the ‘FindMultiModalNeighbors’ function. Then, we used the ‘RunSPCA’ function to perform supervised PCA (sPCA) of the antibody data in the reference atlas and determined anchors in our dataset using the ‘FindTransferAnchors’ function. Lastly, we used the ‘MapQuery’ function of Seurat to transfer the cell type label from the reference atlas to our dataset. For this task, we summarized pre-, pro-, and pre-pro-B cells into one class which we termed pre-B cells. After joining the reference atlas and our data using the sPCA dimension reduction, we generated a new low-dimensional representation of the combined dataset using the ‘RunUMAP’ function.

For associating DNAm differences with gene expression changes, we leveraged the labels transferred from the CITE-seq atlas and correlated the mean expression from the whole transcriptome analysis of Triana *et al*. [11] with the mean DNAm per cell type for the five cell-type labels transferred. Importantly, we only considered those genes that are located closer than 25kb in both orientations from the investigated CpG. Then, we investigated those genes showing the strongest (negative/positive) Pearson correlation with the target CpG. We found that there were more negatively correlated than positively correlated genes and that the CpGs with the strongest correlation were preferentially located in the gene promoter (Suppl. Tab. 7).

## Data visualization

All analyses were performed with R-version newer than 4.0 [12] and the ggplot2 (https://ggplot2.tidyverse.org) and ComplexHeatmap [13] R packages were used for plotting. The lines in the boxplot represent the median, the 25th- and 75th-percentiles, and 1.5 times the inter-quartile range.

