## Supplementary Figures 1-10 for "scTAM-seq enables targeted high-confidence analysis of DNA methylation in single cells"

### Supplementary Figures for "scTAM-seq enables targeted high-confidence analysis of DNA methylation in single cells"

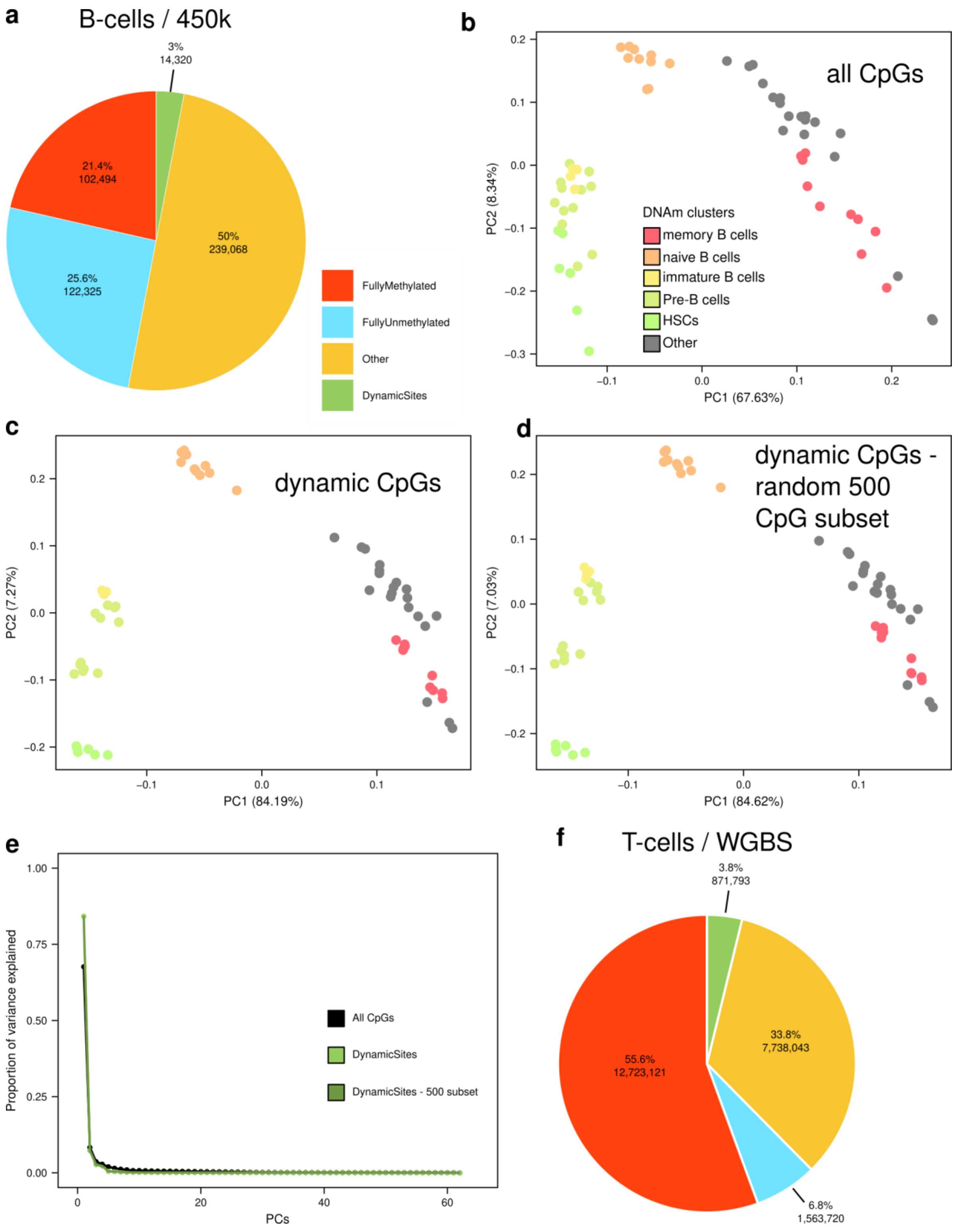


**Supplementary Figure 1:** CpGs dynamically methylated across B- and T-cell development. **a:** Pie chart estimating the number of potentially interesting CpGs in human B-cells (450k data, Kulis et al). We used all the replicates of the different cell types (HSCs, preB, immature, naïve and memory B cells). Fully methylated CpGs are defined as CpGs with methylation value higher than 0.75 in all samples and fully unmethylated are those with less than 0.25. We defined as dynamic sites CpGs showing a DNA methylation difference of at least 0.5 in at least one of the pairwise comparisons between the cell types. All remaining CpGs are classified as 'Others'. PCA for all (478,207) CpGs (**b**), only for the dynamic CpGs (**c**), and for a 500 CpGs randomly selected from the set of dynamic CpGs (**d**). **e:** Proportion of variance explained for the PCs shown in **b, c** and **d**. **f:** Analysis similar to **a** for human T cells (naïve, central-memory and effector-memory T cells, Farlik et al). DNAm data was generated by WGBS and we focused only on those CpGs with at least 10 sequencing reads in all the samples.

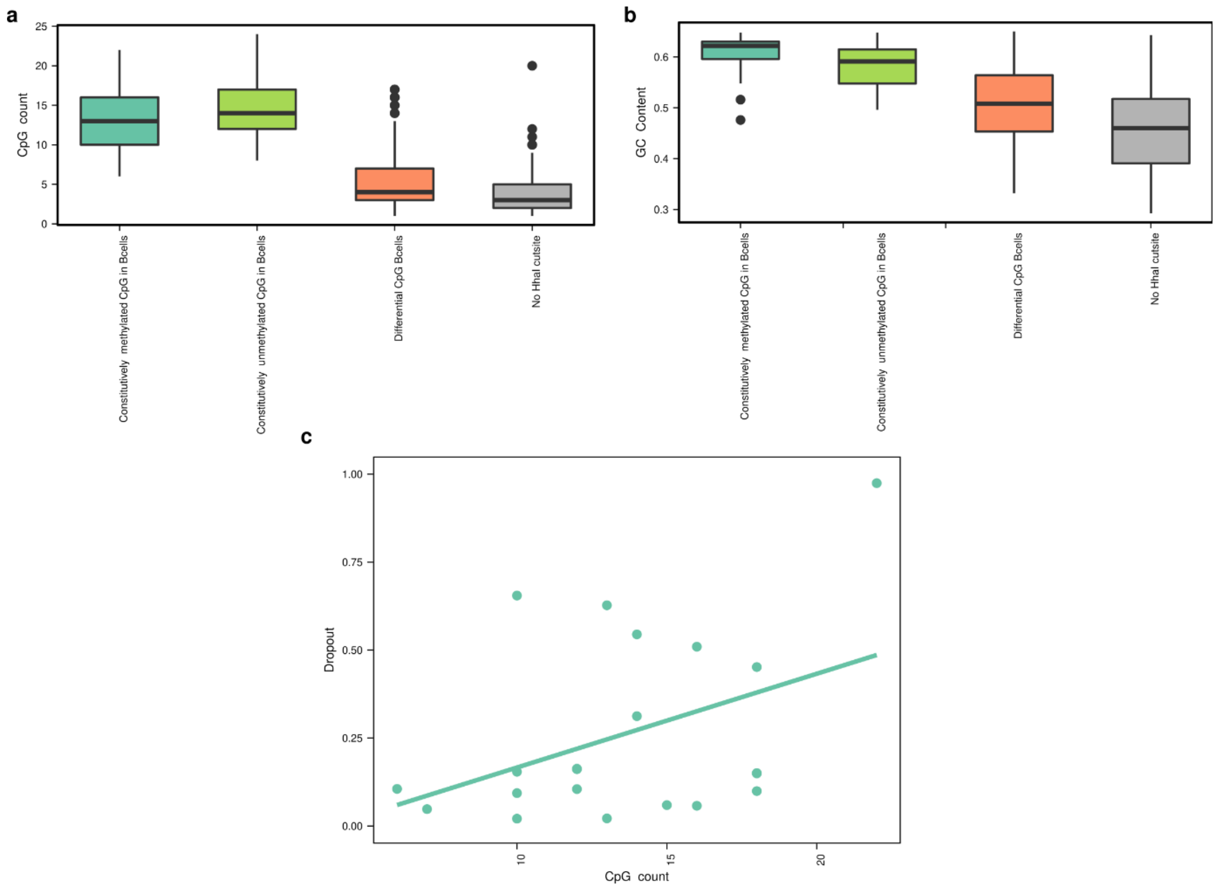


**Supplementary Figure 2:** (**a**) CpG count and (**b**) GC content stratified according to the different amplicon classes. **c**. Relationship between CpG count and dropout rate in the bone marrow sample for the amplicons of class 'constitutively methylated'. The solid line represents the least-squares linear regression line.

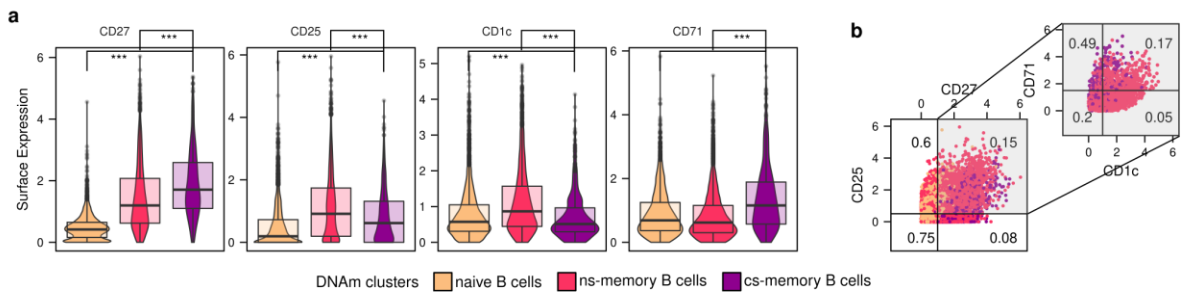


**Supplementary Figure 3:** Surface protein expression in the peripheral blood sample. **a.** Boxplots with single-cell surface protein expression levels. The markers were selected due to their expression profile in previous CITE-seq atlases (Triana et al, 2021). ***: Two-sided Wilcoxon test p-value<0.001. **b.** Scatterplot comparing cell-surface protein expression across different clusters. Numbers in quadrants represent the fraction of naive B cells (left panel) and cs-memory B cells (right panel) across all cells in the quadrant.

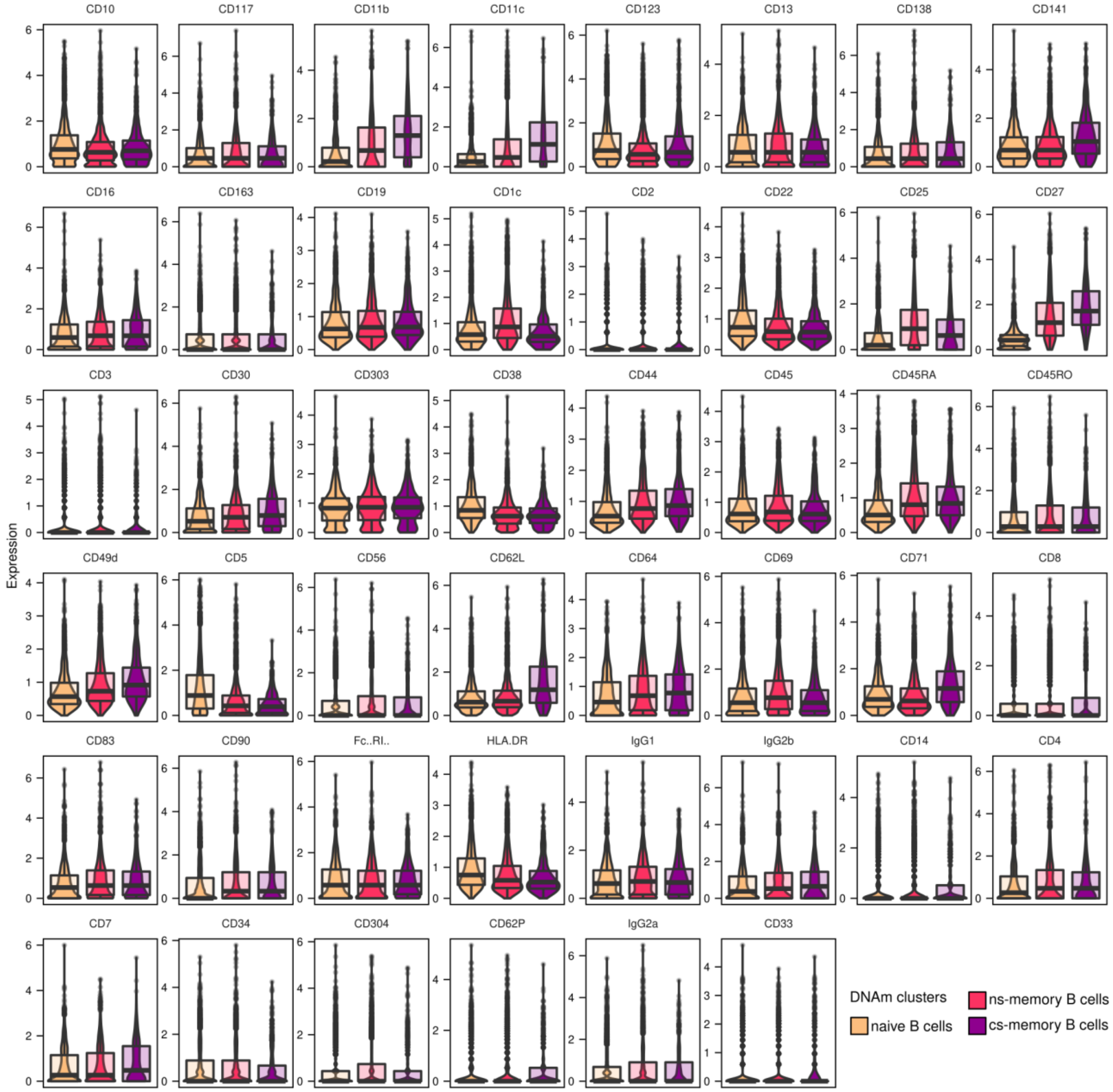


**Supplementary Figure 4:** Combined box- and violin-plot displaying the CLR-normalized expression for all 46 surface proteins analyzed using scTAM-seq across the three DNAm-based cell clusters.

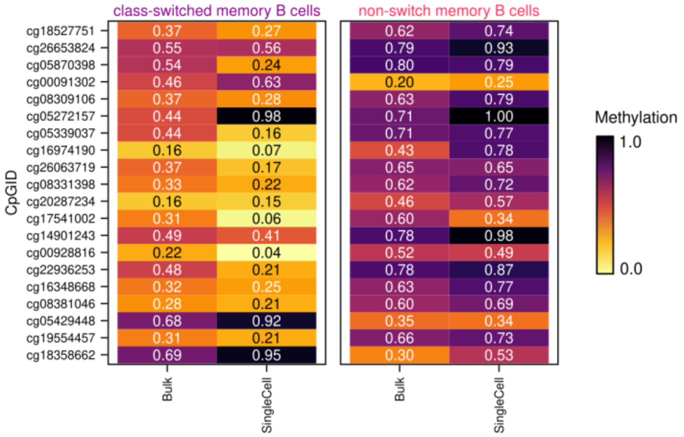


**Supplementary Figure 5:** Comparison of bulk and pseudobulk DNAm values for class-switched (cs-) and non-switched (ns-) memory B cells for the 20 CpGs with the highest mean difference in the bulk data.

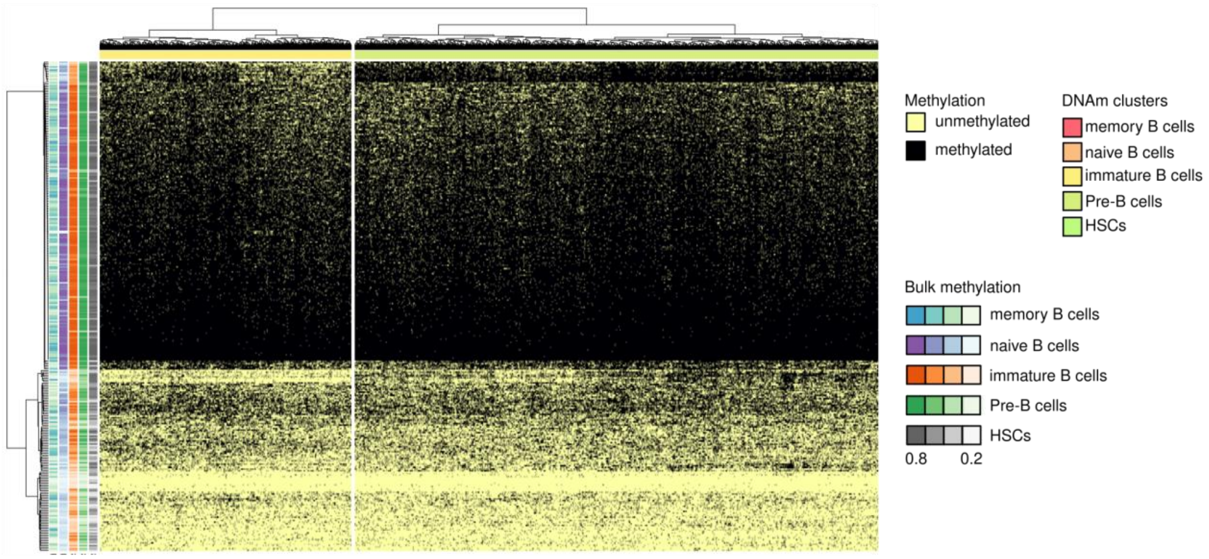


**Supplementary Figure 6:** Heatmap visualizing the DNA methylation states determined using scTAM-seq for the 2,366 cells in the pre/immature B-cell cluster across the 313 selected amplicons.

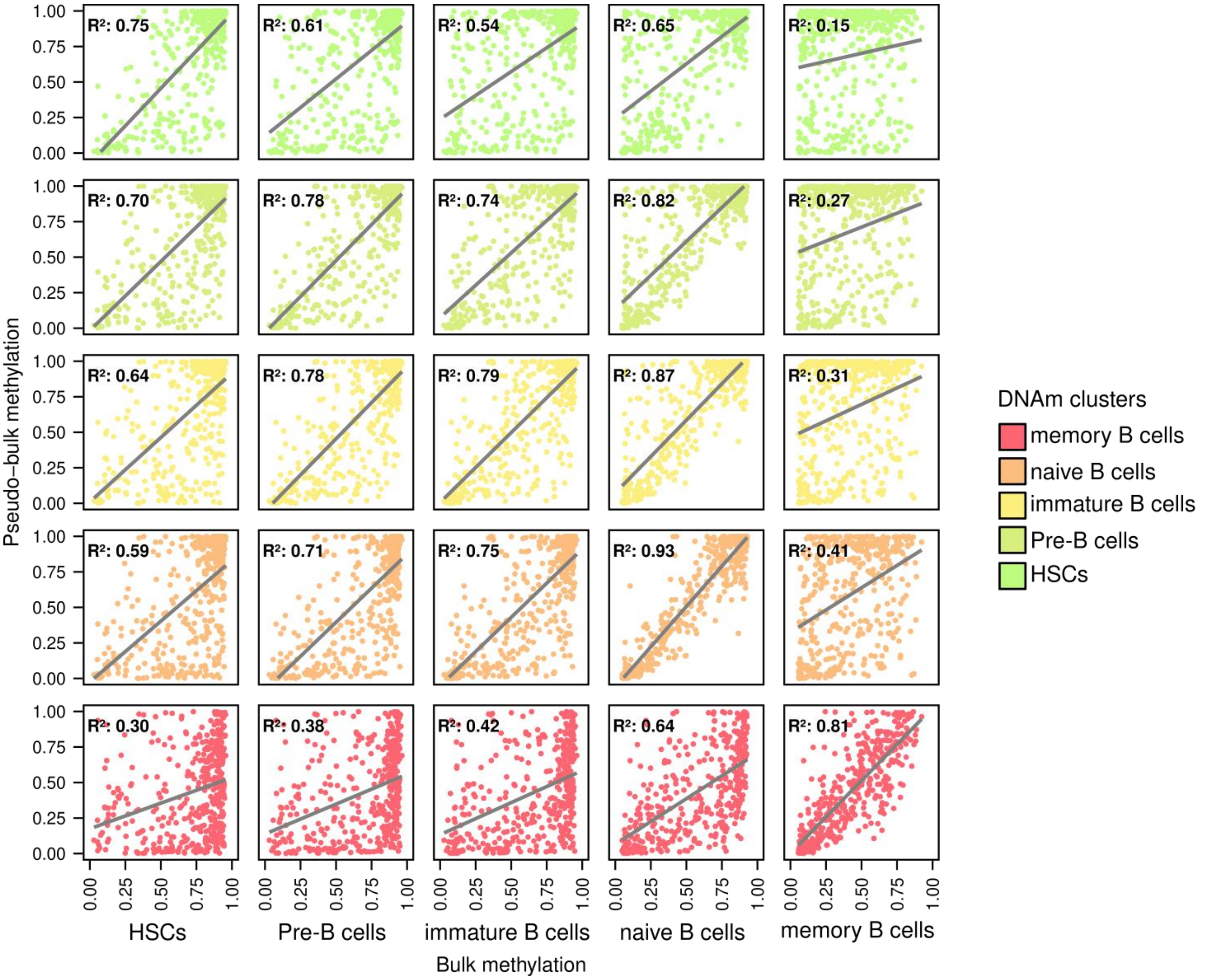


**Supplementary Figure 7:** Comparison of bulk (x-axis) and pseudobulk (y-axis) DNAm value for the five clusters defined on the bone marrow sample for all 424 B-cell-related amplicons. The $R^2$ indicates the Pearson correlation coefficient between x- and y-values and the solid line represents the least-squared regression line.

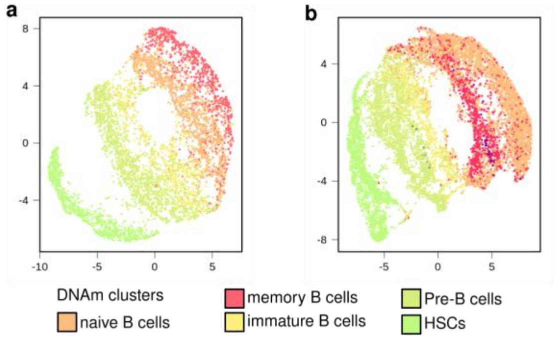


**Supplementary Figure 8:** Visualization of the surface protein data for the bone marrow sample with the cells colored by the cell type labels inferred from the DNAm data. Low-dimensional representation of the cell-surface protein expression as a UMAP (**a**). The surface protein expression data assayed using scTAM-seq was processed using Seurat (see Methods). **b**. Integration with cell-surface proteins data from the CITE-seq reference atlas (Triana et al., 2021). Since scTAM-seq and the reference atlas share 33 antibodies, multi-modal nearest neighbors can be used to generate a joint low-dimensional representation of the two datasets (see Methods).

**Supplementary Figure 9:** Bioanalyzer profiles of libraries from bone marrow digested (**a**, **b**) and undigested (**c**, **d**) samples. **a**, **c**. ScTAM-seq libraries, target peak: 460 bp. **b, d**. Surface proteins

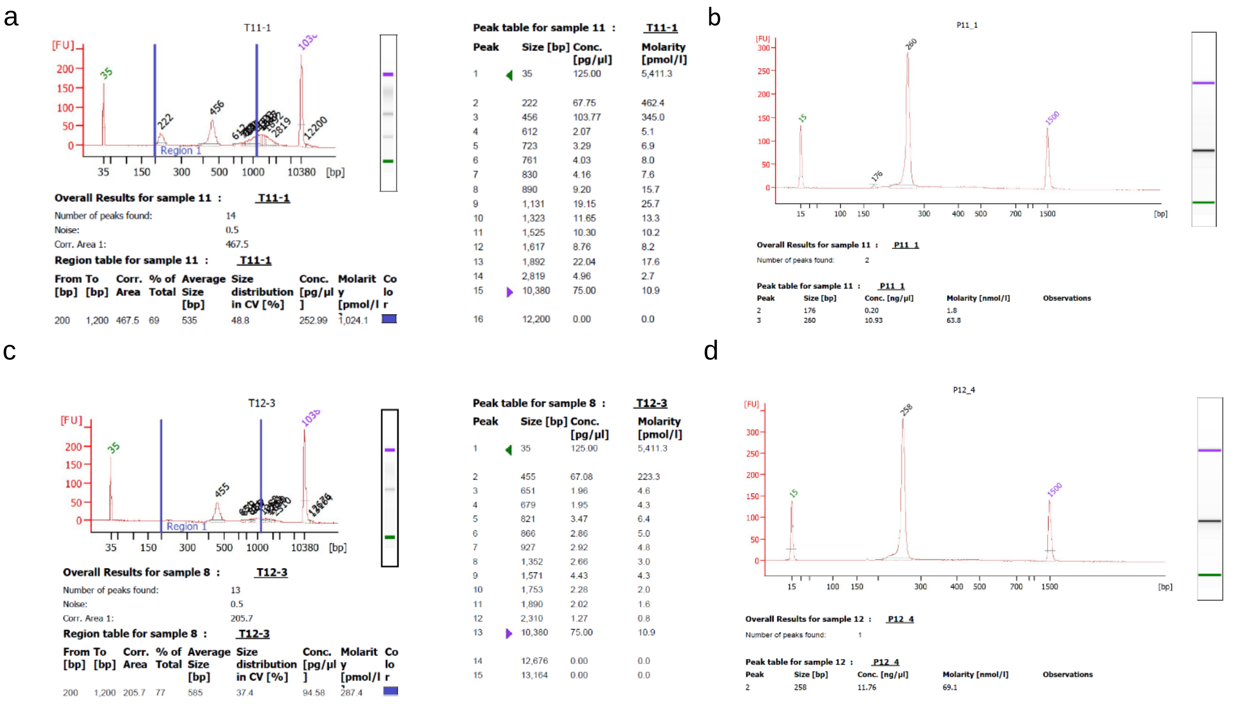


libraries, target peak: 260bp.

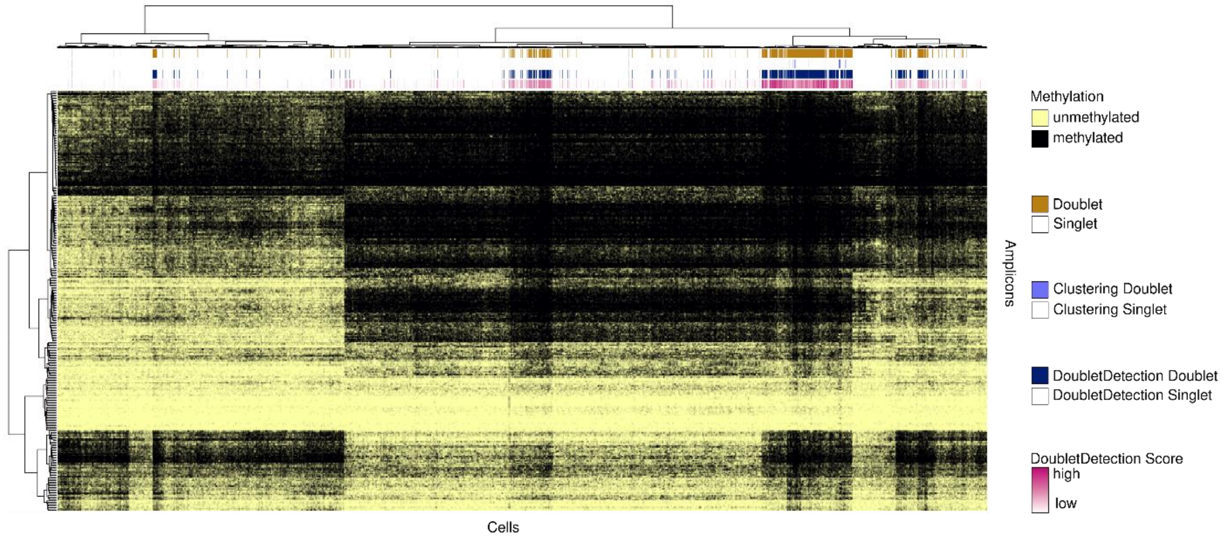


**Supplementary Figure 10:** Doublets as detected by the DoubletDetection software marked in the scTAM-seq heatmap for the digested blood sample. Out of the 11,438 cells, 1,794 cells were marked as doublets by DoubletDetection. Additionally, we identified 61 potential doublets in a per-cluster analysis based on the number of features per cluster (Clustering doublets).
